# Inhibiting the mitochondrial pyruvate carrier does not ameliorate synucleinopathy in the absence of inflammation or metabolic deficits

**DOI:** 10.1101/2020.10.01.322115

**Authors:** Wouter Peelaerts, Liza Bergkvist, Sonia George, Michaela Johnson, Lindsay Meyerdirk, Emily Ku, Jennifer A. Steiner, Zachary Madaj, Jiyan Ma, Katelyn Becker, K. Peter R. Nilsson, Jerry Colca, Patrik Brundin

## Abstract

Epidemiological studies suggest a link between type-2 diabetes and Parkinson’s disease (PD) risk. Treatment of type-2 diabetes with insulin sensitizing drugs lowers the risk of PD. We previously showed that the insulin sensitizing drug, MSDC-0160, ameliorates pathogenesis in some animal models of PD. MSDC-0160 reversibly binds the mitochondrial pyruvate carrier (MPC) protein complex, which has an anti-inflammatory effect and restores metabolic deficits. Since PD is characterized by the deposition of α-synuclein (αSyn), we hypothesized that inhibiting the MPC might directly inhibit αSyn aggregation *in vivo* in mammals. To answer if modulation of MPC can reduce the development of αSyn assemblies, and reduce neurodegeneration, we treated two chronic and progressive mouse models; a viral vector-based αSyn overexpressing model and a pre-formed fibril (PFF) αSyn seeding model with MSDC-0160. These two models present with distinct types of αSyn pathology but lack inflammatory or autophagy deficits. Contrary to our hypothesis, we found that a modulation of MPC in these models did not reduce the accumulation αSyn aggregates or mitigate neurotoxicity. Instead, MSDC-0160 changed the post-translational modification and aggregation features of the αSyn. These results are consistent with the lack of a direct effect on MPC modulation on synuclein clearance in these models.

## Introduction

Synucleinopathies are age-related neurodegenerative diseases characterized by α-synuclein (αSyn) protein misfolding, inflammation and metabolic deficits. People with Parkinson’s disease (PD), dementia with Lewy bodies (DLB) and multiple system atrophy (MSA) present with peripheral and central synucleinopathies that coincide with a wide range of non-motor (e.g., hyposmia, constipation and rapid eye movement sleep behavior disorder) and classical motor symptoms^1^. The disease mechanisms underlying synucleinopathies are not completely understood, but evidence suggests that mitochondrial dysfunction, neuroinflammation and altered protein homeostasis all play a role, as reviewed in detail elsewhere^2^.

Epidemiological studies have indicated a link between PD and type 2 diabetes^3–5^. Type 2 diabetes is associated with a 38% increased risk of developing PD^3,4^. On a molecular level, both people with PD and those with type 2 diabetes have similar metabolic abnormalities such as mitochondrial dysfunction and insulin resistance^5^. Given these links between PD and metabolic dysfunction, clinical trials in PD were initiated to test drugs already approved for the treatment of diabetes. In a randomized, double-blinded placebo-controlled trial with the brain penetrant insulin sensitizing glucagon-like peptide 1 (GLP1)-agonist exenatide, it was shown that exenatide treatment was associated with a reduced decline in motor dysfunction in PD patients^6^. Specifically, once-weekly injections of the slow release form of exenatide for 48 weeks resulted in improvements in the MDS-UPDRS (part III)-defined primary endpoint with effects persisting after a 12-week wash out period^6^. In a post hoc analysis evaluating non-motor symptoms, secondary end points such as patient and observer-led scales assessing mood also showed positive effects from the treatment^7^. Exenatide is currently in a Phase III trial in 200 PD patients who will be treated for 2 years, with the primary endpoint focused on disease modification (ClinicalTrials.gov identifier NCT0423296).

A second insulin sensitizer that has been associated with a lower risk of PD is pioglitazone^3,4^. However, when evaluated in a phase 2 clinical PD trial, pioglitazone did not have any significant effect on disease progression as measured via MSD-UPDRS (part III)^8^. Pioglitazone belongs to a class of anti-diabetic drugs, termed thiazolidinediones or TZDs, and was believed to exert its antidiabetic effects via activation of the transcription factor peroxisome proliferator-activated receptor-γ (PPARγ). Due to strong binding of pioglitazone to PPARγ the drug has several negative side effects. As a result, PPARγ–sparing TZDs, including MSDC-0160 were developed. Interestingly, when compared to pioglitazone, MSDC-0160 has similar anti-diabetic effects but it does not activate PPARγ, indicating that TZDs act on a different receptor to elicit effects. The primary target was later found on the inner membrane of mitochondria and was identified as the previously unknown mitochondrial pyruvate carrier (MPC)^9^.

Because of the link between TZDs and a lowered associated risk of PD, MSDC-0160’s limited side effects and its bioavailability in brain, we previously tested it in cell and animals models of PD and found that MSDC-0160 is neuroprotective^10^. Specifically, MSDC-0160 protects against dopaminergic cell loss in the substantia nigra in neurotoxin-challenged mice and in a genetic mouse model with hemizygous loss of the transcription factor Engrailed 1 (En1), which is believed to entail mitochondrial deficits^11^. We also observed a mitigation of neuron death of MSDC-0160 treatment in *Caenorhabditis elegans* (C. elegans) in which we over-expressed αSyn. Taken together, we hypothesized that the beneficial effects of MSDC-0160 were due to its antiinflammatory effects and its ability to normalize metabolic deficits.

The effects of MSDC-0160 on αSyn aggregation in progressive, chronic PD models have previously not been investigated. In this study, we therefore evaluated MSDC-0160 in two different α-syn based rodent models of PD; 1) a rat model with adeno-associated virus (AAV) vector over-expression of human αSyn and 2) a mouse αSyn pre-formed fibril (PFF) seeding model. Unexpectedly, we found that MSDC-0160 increased the levels of aggregated αSyn in the AAV-overexpression model, possibly via increased oxidation of soluble αSyn. In the PFF seeding model, MSDC-0160 also increased αsyn burden transiently; a significant difference between the treatment groups was observed five weeks after PFF injection, but it was not present at the later 13-week timepoint. We examined both models further examined for inflammatory and lysosomal markers, and found them both to be unaltered, even in the presence of robust αSyn pathology. We conclude that while neuroprotective actions of MSDC-0160 may occur via restoring inflammatory and metabolic deficits, we could not demonstrate effects on protein aggregation or protein spreading in these rodent models of synucleinopathies.

## Methods

### Animals

Twelve-week-old male and female C57BL/6J wildtype (WT) mice were sourced from The Jackson Laboratory. Eight-week old female rats were obtained from Charles River. Animals were housed with a maximum of four mice or two rats per cage under 12-h light/12-h dark cycles with free access to food and water. Mice and rats were fed a diet of chow formulated to deliver MSDC-0160 (30 mg/kg) or control chow starting at 1 week after stereotactic surgery. Mice were euthanized at two different time points: one and three months after PBS/PFF injections. Rat were euthanized four months post viral transduction. The housing of animals and all procedures were performed in accordance with the *Guide for the Care and Use of Laboratory Animals* (United States National Institutes of Health) and were approved by the Van Andel Research Institute’s Animal Care and Use Committee.

### PFF production and OB injections

Mouse αSyn aggregates were produced as described previously^12^. Before surgery, PFFs were prepared by the sonication of αSyn aggregates in a water-bath cup-horn sonicator for four min (QSonica, Q700 sonicator, 50% power, 120 pulses 1 s ON, 1 s OFF). Mice were anesthetized with isoflurane/oxygen and injected unilaterally in the right olfactory bulb with either 0.8 μL of PFF (5 μg/μl; n=36, 18 females/males) or 0.8 μL of PBS as a control (n=36, 18 females/males). Coordinates from bregma: AP: + 5.4 mm; ML: +/- 0.75 mm and DV: - 1.0 mm from dura. Injections were made at a rate of 0.2 μL/min using a glass capillary attached to a 10 μL Hamilton syringe. After injection, the capillary was left in place for three min, before being slowly removed.

### Viral vector production and nigral injections

rAAV2/5-CMVSyn-human-αSyn and rAAV2/5-CMVSyn-GFP production was performed as described previously^13^. Genome copies were determined via qPCR and two viral vector titers were used for injection into the rat SN. Nigral injections were performed under general anesthesia using an isoflurane/oxygen mixture. For the αSyn low dose and αSyn high dose a total of 1.2×10^11^GC and 2.5×10^11^GC were injected, respectively. As a control, 3×10^11^GC of GFP expressing viral vector or PBS were injected. Coordinates from bregma: AP: 5.3 mm, ML: 2.0 mm and DV: 7.2 mm measured from dura^14^. A volume of 3 μl of viral vector was infused at a rate of 0.25 μl/min using a 30G needle and 10 μl Hamilton syringe.

### MSDC-0160 metabolite analysis

Mice and rats were allowed free access to chow containing either MSDC-0160 (30 mg/kg) or placebo for four months. For mice, whole blood was collected one and three months after PFF injection from the submandibular vein in 6 mL BD heparinized tubes. For rats, whole blood was collected in identical tubes via cardiac puncture four months after viral vector injection. The samples were centrifuged for 15 min at 4°C (2000 x g), followed by collection of the supernatant (plasma) which was snap frozen and stored at −80°C.

Frozen heparinized plasma samples were blind-coded and analyzed by Charles River Laboratories (Mattawan, MI) for the concentrations of the MSDC-0160 metabolite MSDC-0037 in Study Protocol 1443-007B. In short, each 25 μL aliquot of standard, QC sample, or study sample was mixed with 10 μL of working internal standard solution (1,250 ng/mL in methanol/water [50/50, v/v]) and 140 μL of water. The samples were vortexed and transferred to an ISOLUTE SLE plate and eluted with 1.0 mL of MTBE. The samples were evaporated and reconstituted with 100 μL of methanol followed by 100 μL of 5 mM ammonium formate in water. The samples were mixed and transferred to a clean 96-well plate. An aliquot was injected onto an LC-MS/MS system for analysis. The liquid chromatography system used a MacMod Ace 3 C18 column, 2.1 x 50 mm (3 μm particle size) with a gradient flow consisting of 5 mM ammonium formate in water/methanol (75/25, v/v) and 5 mM ammonium formate in methanol/acetonitrile/water (72/18/10, v/v/v) at a flow rate of 500 μL/minute.

The analyte, metabolite, and internal standard were detected using a SCIEX API 5000 triple quadrupole LC-MS/MS system equipped with an ESI (TurboIonSpray®) ionization source operated in the positive ion mode. The multiple reaction monitoring transitions of the respective [M+H]^+^ ions were used to monitor MSDC-0160 and MSDC-0037 as have previously been described for clinical studies^15^.

### Western blotting

Rats were euthanized with sodium pentobarbital (130 mg/kg; Sigma) and transcardially perfused with PBS to remove blood. Brains were isolated and the substantia nigra was dissected on ice using a rat brain matrix. Isolated samples were snap frozen and stored at −80 °C for later analysis. Frozen tissue was weighed in equilibrated Eppendorf tubes and homogenized in PBS 10% w/v with protease and phosphatase inhibitors (ThermoFisher). Homogenization was performed by by probe sonication at 4 °C for 2 rounds of 15 s pulses at 0.5 Hz with a 10% amplitude. The whole homogenates were centrifuged for 10 min at 6000 x g at 4 °C. The pellet was discarded and supernatant was collected for analysis. To isolate insoluble αSyn, 20% sarkosyl was added to the whole homogenate to result in a 1% sarkosyl PBS sample that was incubated on a rotating shaker for 1 h at room temperature. After incubation, samples were centrifuged for 10 min at 6000 x g at 4°C to remove remaining cell debris. The cleared supernatant was centrifuged at 100,000 x g for 60 min and the resulting pellet was gently washed in PBS after which it was resuspended in 1% sarkosyl in PBS with protease and phosphatase inhibitors. A second centrifugation step at 100,000 x g for 10 min was performed to collect the 1% sarkosyl insoluble pellet. Protein concentrations were estimated using a BCA kit (Thermo Fisher). Samples were prepared with Laemmli buffer containing 2% SDS, heated at 95°C for 10 min for denaturation and stored at −80°C. Protein samples were separated via 4-15% SDS PAGE (BioRad) and transferred to a PVDF membrane using the BioRad Turboblot system. PVDF membranes were blocked with 5% BSA in PBS during 30 min at room temperature. Overnight incubation with primary antibody, Iba-1 (1:500, WAKO), human αSyn (4B12, 1:1000, BioLegend), pSer129-αSyn (pS129, 1:5000, Abcam), LAMP1 (1:1000, Abcam), caspase (1:500, Abcam) was followed by incubation with HRP-conjugated secondary antibody (Cell Signaling Technology). Signal was detected by chemiluminescence (Pico Chemiluminescent Substrate, Thermo Fisher) using a BioRad Imager. Western blot bands were quantified using ImageJ software.

### Behavioral analysis

To monitor functional deficits related to nigral dopamine neuron dysfunction and death, rats were subjected to a cylinder test to evaluate spontaneous forelimb use. Rats were placed in a clear glass cylinder and were filmed for a total of 30 contacts with the cylinder. Percentage of forepaw use was expressed as the percentage of right forepaw touches over total number of touches. Non-lesioned rats score around 50%.

To examine olfactory function in mice, the buried pellet test was carried out as recently described by our laboratory in Johnson et al.^16^. In short, the mice were fasted overnight starting three days prior to testing. Throughout the experiment, their body weight was monitored and mice who lost more than 10% of their original body weight were excluded from further fasting and from the experiment. At the first day of testing, a surface test was performed, where the treat (Bio-Serv, Fruit Crunchies) was placed on top of the clean bedding. Mice who were not motivated to eat the treat were excluded from further testing (all mice in this study were motivated by the treat and were thus included). For four consecutive days, the treat was buried at different locations 1 cm under the clean bedding and latency for mice to uncover it was recorded. A maximum time was set at 300 s, and if a mouse did not uncover the treat within that time frame, 300 s was recorded as its latency. The average latency to uncover the treat per animal is presented here, with nine to ten animals analyzed per group.

### Immunohistochemistry

Mice and rats were euthanized with sodium pentobarbital (60 mg/mL; Sigma) and transcardially perfused with room temperature 0.9% saline followed by ice-cold 4% paraformaldehyde (PFA) in 0.1 M phosphate buffer. Brains were removed, post-fixed overnight at 4°C in 4% PFA and subsequently placed in 30% sucrose. Brains were frozen and coronal sections of 40 μm were cut on a microtome (Leica) and collected as serial tissue sections. For immunohistochemistry in mice, a series (every 240 μm) of coronal free-floating sections were stained using 1:10000 anti-pSer129 αSyn primary antibody (Abcam, Ab51253), 1:1000 anti-NeuN primary antibody (Millipore, Ab377), 1:5000 anti-nitrated αSyn primary antibody (Thermo Fisher, 35-8300) or 1:800 anti-Iba1 primary antibody (Wako, 019-19741) over night. For immunohistochemistry in rats, a series (every 240 μm) of coronal free-floating sections of SN was stained using 1:10000 anti-pSer129 αSyn primary antibody (Abcam, Ab51253), 1:5000 anti-nitrated αSyn primary antibody (Thermo Fisher, 35-8300), 1:2000 anti-TH (Abcam), or 1:800 anti-Iba1 primary antibody (Wako, 019-19741) overnight. Corresponding secondary antibodies (1:500 goat anti-rabbit/goat anti-mouse biotinylated secondary antibody (Vector Laboratories, BA-1000 and BA-9200) were added. The antibody signal was amplified using a standard peroxidase-based method (VectaStain ABC kit) and developed using a DAB kit (Vector Laboratories). Stained tissue sections were mounted onto gelatin coated glass slides. NeuN- and TH-immunostained slides were counter stained with cresyl violet (CV). After dehydration, slides were coverslipped with Cytoseal 60 mounting medium (Thermo Fisher Scientific).

For LAMP2 immuostaining, using fluorescence to detect antibody binding, sections were incubated overnight with 1:1000 anti-LAMP2 primary antibody (Abcam, Ab13524) overnight, followed by a 2 h incubation with a fluorescent secondary antibody (Thermo Fisher/Invitrogen, Alexa Fluor 488). DAPI was used to visualize cell nuclei. For the co-stain between h-FTAA and pSer129, tissue sections were incubated with 1:1000 anti-pSer129 αSyn primary antibody (Abcam, Ab51253) for 2h at RT followed by a 1h incubation with 1:200 secondary goat anti-rabbit antibody Alexa 647 (Thermo Fisher, A32733) and DAPI to visualize cell nuclei. Finally, 0.5 μM h-FTAA was added and allowed to incubate with the tissue for 30 min at RT. Fluorescently stained tissue was coverslipped using hard-set antifade mounting media (Vector Laboratories, H-1400). A Nikon A1plus-RSi Laser Scanning Confocal Microscope (Nikon) was used to image these sections.

### pSer129 quantification

Z-stacks of mounted αSyn pSer129-stained tissue sections were captured at 20x magnification using a whole slide scanner (Zeiss, Axioscan Z1) at a 0.22 μm/pixel resolution. Extended depth focus (EDF) was used to collapse the z-stacks into 2D images as they were collected. Tissue thickness was set to 20 μm, z-stacks were collected at 3 μm intervals and the method used was Contrast. The images were exported with 85% compression as .jpeg files. The digitized images were then uploaded Aiforia™ image processing and management platform (Aiforia Technologies, Helsinki, Finland) for analysis with deep learning convolution neural networks (CNNs) and supervised learning. A supervised, multi-layered, CNN was trained on annotations from digitized αSyn pSer129-stained coronal slices to recognize total αSyn-positive staining. The algorithm was trained on the most diverse and representative images from across multiple pSer129 datasets to create a generalizable AI model capable to accurately detecting αSyn pSer129-stained profiles in slides collected by multiple investigators. Ninety-one images constituted training data. The ground truth, or features of interest used to train the AI model, were annotated for each layer within the Aiforia™ Cloud platform and constituted input data for each CNN. The first feature layer was annotated using semantic segmentation to distinguish the total tissue from the glass slide. The second feature layer was annotated using semantic segmentation to distinguish total αSyn pSer129-positive staining. Features that were considered artifact (glass slide, debris) were annotated as background, and constituted additional input training data for the multi-layered CNN. The regions corresponding to the AON and the PRh were outlined according to the Allen Mouse Brain Atlas (Allen Institute) and the CNN was used to quantify the area percentage that contained αSyn pSer129-positive signal. For each brain region, the average area with αSyn pSer129-staining is reported as a percentage of the whole area assayed. For the one-month time point, six animals per group were analyzed and for the three-month time point seven to eight animals per group were included in the analysis.

### Stereology

Surviving dopamine neurons and total neuronal cell counts were quantified using unbiased sampling and blinded stereology. For the AAV αSyn over-expression model, tyrosine hydroxylase (TH), αSyn pSer129 and nitrated αSyn-positive cells were counted in the SNpc and SNpr in a series of seven sections at an interval of six sections at 40 μm thickness (240 μm intervals). Contours of the regions were drawn at 10x magnification and quantifications were performed at 60x using a 200 μm counting frame. Dissector height was 12 μm with a 3 μm guard zone. The Gunderson estimated m=1 error was less than 0.1 and a total of eight animals per group was counted. For the αSyn PFF seeding model, NeuN- and CV-positive cells were counted in the AON, which was outlined according to the Allen Mouse Brain Atlas (Allen Institute) using a 2.5x objective. The quantification of NeuN and CV-positive cells was performed on every 6^th^ AON (240 μm) section using the optical fractionator probe in Stereo Investigator software (MBF Bioscience). The counting was accomplished using a 60x oil objective, with the grid set to 200×200 and the frame set to 40×40. The dissector height used was 10 μm, with a 3 μm guard zone. The Gunderson estimated m=1 error was less than 0.1 and a total of nine to ten animals per group were counted.

### Assessment of microglia morphology

Microglia hydraulic radius (area/perimeter ratio) was investigated as previously described ^13^. In short, color (RBG) images were generated using a 60x oil objective. Following imaging, a custom MATLAB script assessed the morphology of the imaged microglia, calculating the area/perimeter ratio. For the PFF seeding model, five to six animals per group were analyzed. Microglia was imaged from every 6^th^ AON section and a minimum of 30 individual microglia was analyzed per animal, with the average microglia area/perimeter ratio reported.

### Quantification of fluorescent LAMP2 signal

Images corresponding to the AON were captured using a 20x oil objective on a Nikon A1plus-RSi Laser Scanning Confocal Microscope (Nikon) in a blinded manner. The area percentage occupied by LAMP2 positive signal was quantified using FIJI software (Nation Institute of Health) after an unbiased threshold was applied. The average area percentage calculated for each animal is reported here. For the PFF seeding model, a total of six mice per group were analyzed.

### Statistics

All AAV and PFF mouse model data were analyzed using R v3.6.0 (https://cran.r-project.org/) and assumed two-sided hypothesis tests with a significance level <0.05 after Benjamini-Hochberg multiple testing corrections. In the PFF model, neuronal cell count, Iba1 and pSer129 quantification were done in parallel with another treatment group (PT302, data not shown), meaning they share the same set of placebo treated animals; the multiple testing corrections take into these additional comparisons. For experiments that analyzed both hemispheres of the brain independently (e.g. ipsilateral versus contralateral) the appropriate mixed-effects model was used (R package *lme4* https://cran.r-project.org/web/packages/lme4/index.html unless otherwise noted). Count data were analyzed using negative binomial regressions; mixed-effects negative binomial regressions were done using the R package *glmmTMB* and the ‘nbinom1’ family (https://cran.r-project.org/web/packages/glmmTMB/index.html). Time-to-event data (e.g. the buried pellet test) were analyzed using Cox proportional hazards regression. All other data types were analyzed via linear regression with log and square-root transformed outcomes as needed based on standard regression diagnostics (i.e. normality of residuals and homoscedasticity) (log: IF αSyn pSer129 and soluble αSyn; square-root: soluble αSyn pSer129, insoluble human αSyn, insoluble αSyn pSer129). Cook’s distance was used to assess individual observations with high amounts of leverage (set as 4/(n-p-1)). For linear regressions with highly influential observations, robust regressions with MM estimation were used. Zero variance data (i.e. DAB αSyn pSer129+, IF αSyn pSer129+, soluble human αSyn, and Nissl+ measures across both naive and GFP injected animals), logistic regression with a Firth correction was used to determine if the odds of detection versus no detection differed between groups. An equivalence interval of +/-10% was used to determine if groups had strong evidence for equivalence based on their 95% confidence intervals.

## Results

### Inhibiting the MPC in a viral vector-based model of human α-syn overexpression

To determine the effects of inhibiting the MPC on αSyn aggregation, we used a slowly progressing viral vector-based model that overexpresses αSyn. We selected two doses of viral vector, which from here on will be referred to as ‘αSyn low’ (4.3×10^13^GC/ml) and ‘αSyn high’ (8.6×10^13^GC/ml). These two titers yield a 4- and 6-fold overexpression of αSyn in rat SNpc, respectively (Fig. 1A, 3B).

**Fig 1.**
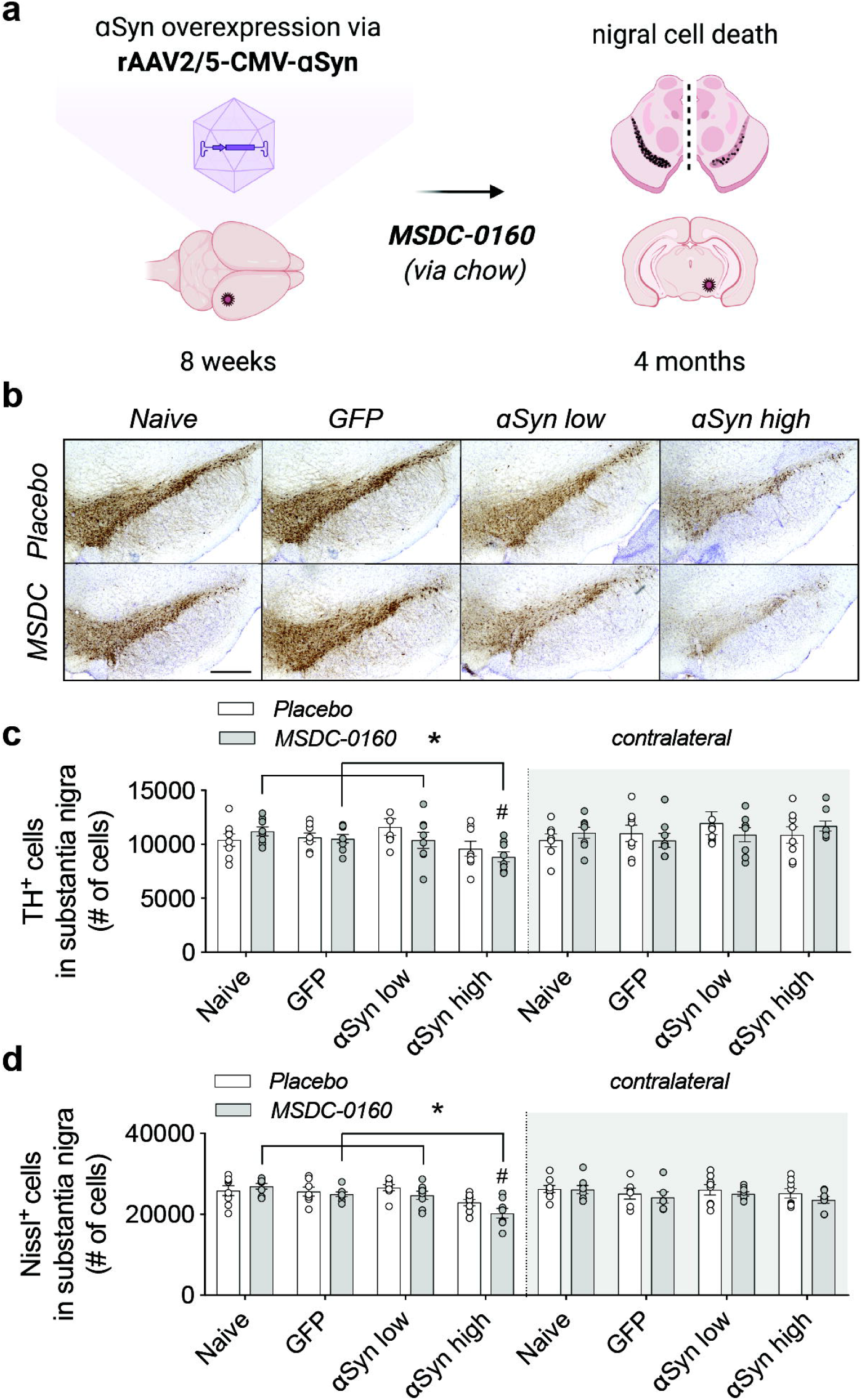
MSDC-0160 treatment does not influence nigral dopaminergic cell viability in a progressive αSyn overexpression model. **a)** Experimental overview of the αSyn viral vector overexpression model in rat SN. **b**) TH expression and Nissl^+^ cells in rat SN for animals treated with placebo or MSDC-0160 in control or experimental conditions. Scale bar = 500 μm. **c)** Stereological quantification of total dopamine neurons in rat SNpc for ipsi- and contralateral sides. The contralateral hemisphere had significantly more TH cells than the ipsilateral in the αSyn high mice (# p = 0.001, ~22.5% more cells CI = 10.3% - 36.0%, negative binomial mixed-effect model, n = 8, s.e.m.). In the ispilateral side, αSyn high had significantly fewer TH cells than Naïve, GFP and αSyn low (*p = 0.031, 14.9% fewer CI = 4.8 – 23.9, *p = 0.05, 13.4% fewer 95% CI 3.1 - 22.6 and *p = 0.0313, 15.6% fewer CI = 24.5-5.6, respectively, negative binomial mixed-effect model, n = 8, s.e.m.). **d**) Stereological quantification of Nissl^+^ cells in rat SN for ipsi- and contralateral sides. For the αSyn high group, the ipsilateral side has ~ 13% fewer Nissl+ cells across both treatment groups than the contralateral (#p < 0.0001, 95% CI 6.4 - 20.1, negative binomial mixed-effect model, n = 8, s.e.m.). αSyn high shows significantly fewer Nissl+ cells compared to animals injected with naive, GFP, or low amounts of αSyn (p < 0.001 for all 3, GFP: 15% fewer CI = 8% - 21%, Naive: 18.4% fewer CI = 12.0% - 24.4%, αSyn low: 16% fewer CI = 22.2% - 9.3%, negative binomial mixed-effect model, n = 8, s.e.m.).

Female rats were stereotactically injected in SNpc at 8 weeks of age with rAAV2/5 CMV-Syn-αSyn or rAAV2/5 CMV-Syn-GFP expressing vector that served as a control (Fig. 1A). One week after injections, the animals were allowed free access to chow containing either MSDC-0160 (30mg/kg) or placebo for four months. During the time of chow administration, we did not observe any differences in animal body weight or chow consumption (Supplementary fig. 1A, B). At the four-month timepoint, plasma was collected and the presence of hydroxymitoglitazone (MSDC-0160 metabolite) was quantified (Supplementary fig. 1C), showing the presence and metabolization of MSDC-0160 at expected levels.

Motor behavior was examined four months after viral transduction using the cylinder test. No difference in motor behavior was apparent between any of the experimental groups, nor did we observe any statistically significant differences between placebo or MSDC-0160 treated groups (Supplementary fig. 2). When assessing the total number of TH- and Nissl-positive neurons in the SN via stereology we find a significant decrease in TH- and Nissl-positive cells between the αSyn high group and all other experimental treatment groups (Fig. 1A-B), suggesting dose-dependent neurotoxic effects of αSyn expression. No cell loss was observed in the GFP group, showing that no toxic effects are due to the viral vector itself. In addition, when the SNpc is separately analyzed for its anterior (Bregma −4.8 mm) and posterior (Bregma −6.2 mm) areas, TH loss in the anterior is much more significant compared to the posterior region (Supplementary fig. 3 A-D). This indicates that αSyn expression from the overexpressing αSyn vector is higher in the anterior SNpc since it is adjacent to the injection site, and lower in the posterior portion, which again confirms vector-driven αSyn neurotoxic effects with no neurotoxicity observed for the GFP control group.

Next, we performed stereological counts of nigral cells positive for phosphorylated αSyn (αSyn pSer129) (Fig. 2A). Unexpectedly, in the αSyn low group, animals treated with MSDC-0160 show significantly more αSyn pSer129+ cells compared to the placebo group (12132 versus 7918 cells, **p = 0.002, negative binomial regression with Benjamini-Hochberg multiple testing adjustments, SEM, n = 8) (Fig. 2B). No significant effect was detected in the αSyn high group between MSDC-0160 and the placebo group.

**Figure 2.**
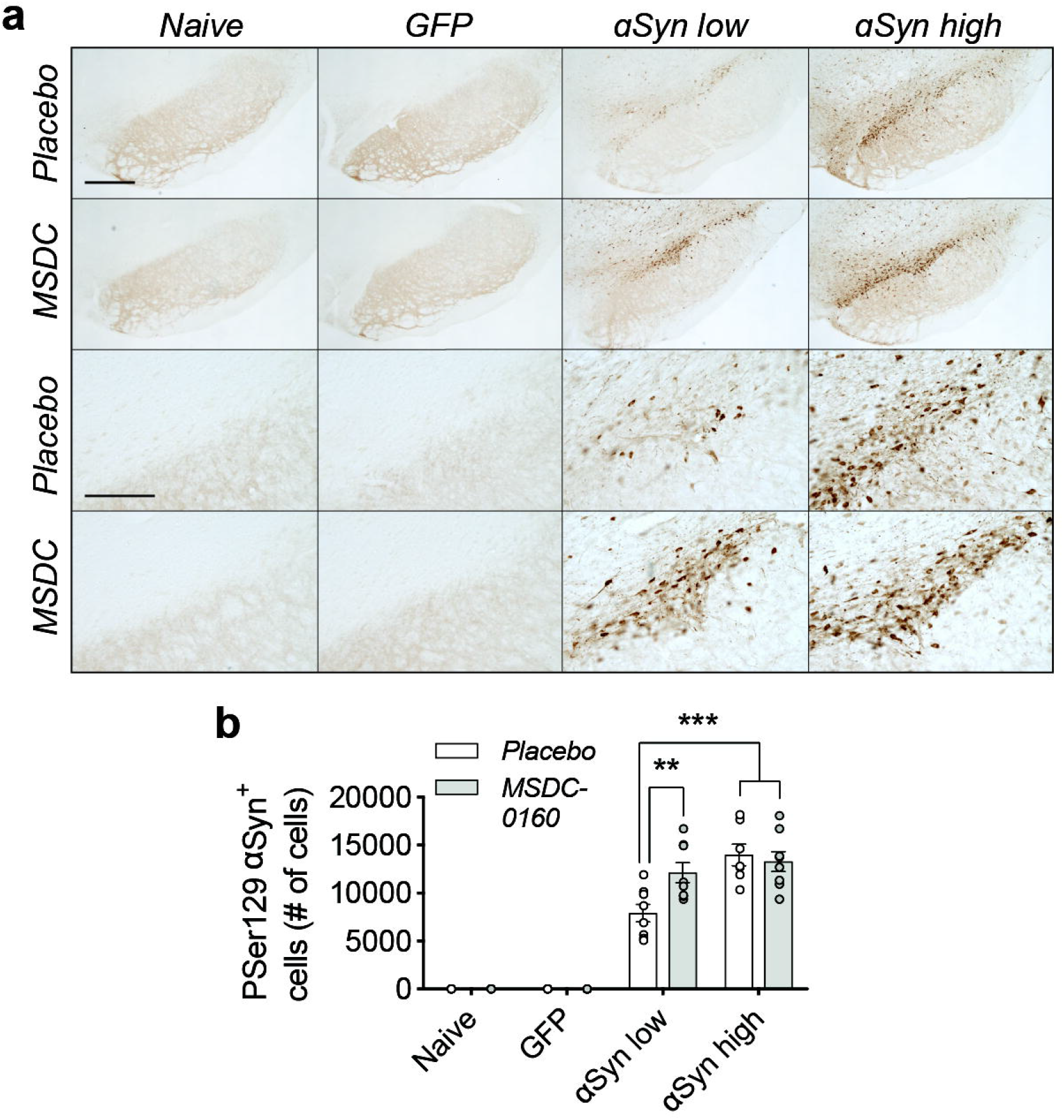
A metabolic switch leads to increased αSyn pathology in rat SN. **a)** Overview of pSer129-αSyn positive cells in rat SN. Scale bars for higher and lower magnification represents 450 μm and 100 μm, respectively. **b)** Stereological quantification of αSyn pSer129 positive cells in rat SN. Treatment with MSDC-0160 leads to a significant increase in phosphorylated cells in the αSyn low group compared to placebo (n = 8, SEM, **p < 0.01 and ***p < 0.001, negative binomial regression with Benjamini-Hochberg multiple testing adjustments, SEM, n = 8)).

We further analyzed phosphorylated levels of αSyn via biochemical analyses and found that both soluble αSyn pSer129 as well as 1% sarkosyl insoluble αSyn pSer129 (1% P129-αSyn) are increased in the MSDC-0160 treated animals compared to its placebo group (*p = 0.046, 95% CI 0.039 – 0.471, square-root linear regression, SEM, n = 6 for soluble αSyn pSer129 and *p = 0.013, 95% CI 0.129 – 0.691, square-root linear regression, SEM, n = 6 for 1% sarkosyl insoluble αSyn pSer129) (Fig. 3E, F). This increase was observed in the αSyn high group between MSDC-0160 and placebo animals whereas a similar trend was noticeable in the αSyn low group between treatment conditions (Fig. 3E, F). This indicates that administration of MSDC-0160 results in increased αSyn aggregation in the slowly progressing viral vector-based model that overexpresses αSyn.

**Figure 3.**
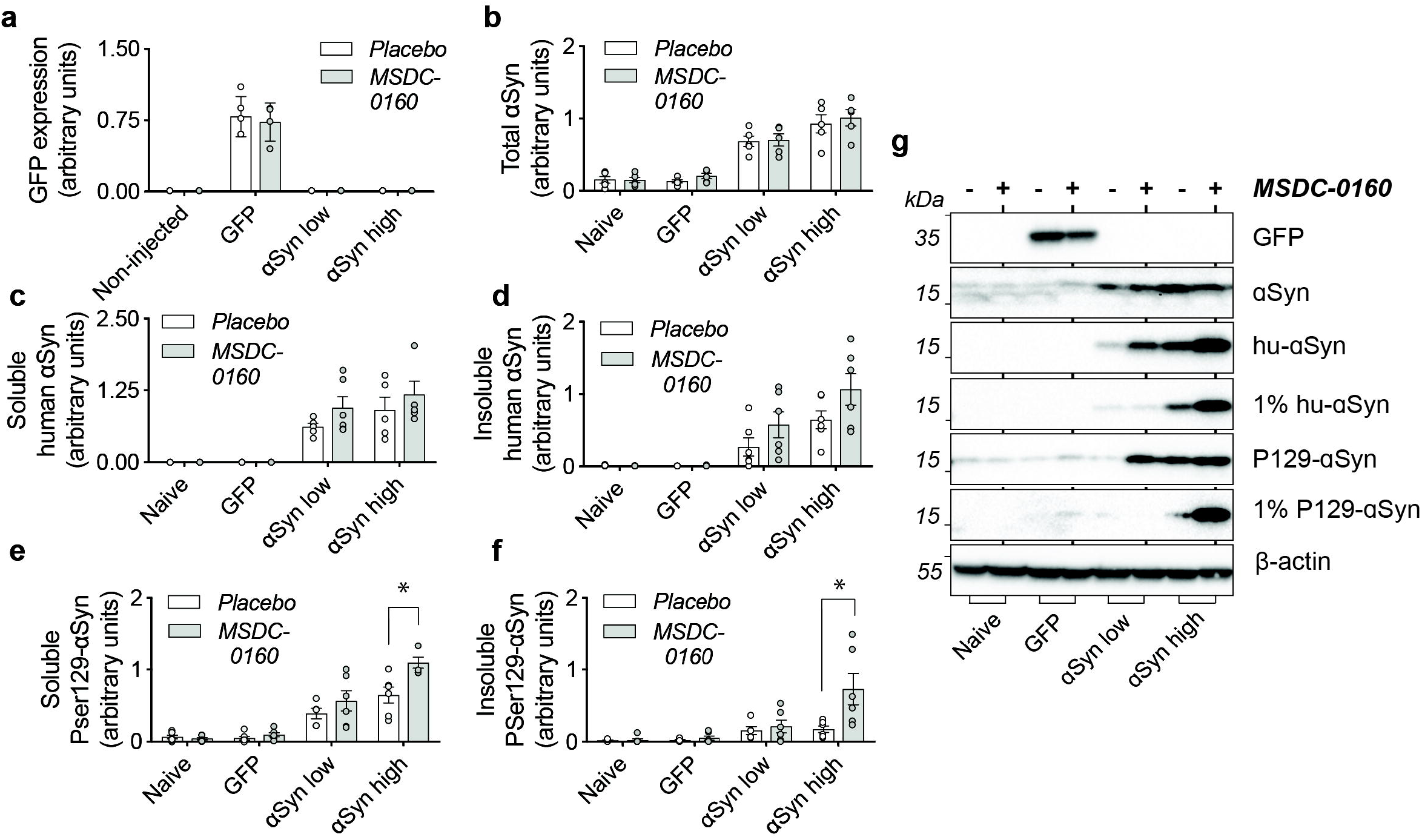
αSyn aggregation is increased in rat SN after inhibition of the MPC. **a)** Inhibition of the MPC has no effect on viral vector-mediated expression of GFP. **b)** Detection of total levels of αSyn in naïve and experimental conditions. αSyn overexpression leads to a 4- and 6-fold increase in αSyn low and high conditions compared to naïve rats. **c,d**) Isolation of sequentially extracted soluble and 1% sarkoysl insoluble human αSyn (1% hu-αSyn) shows no significant differences in aggregated αSyn after treatment with MSDC-0160. **e,f**) Inhibition of MPC leads to significantly increased levels of soluble and 1% sarkosyl insoluble αSyn pSer129 (1% P129-αSyn) in the αSyn high group (n = 6, SEM, *p < 0.05 square-root linear regression, SEM, n = 6). **g)** Representative Western Blot figures for different conditions tested.

αSyn has been shown to bind the extracellular membrane of mitochondria ^17^. Due to its membrane curvature and the presence of the negatively charged cardiolipin in the mitochondrial outer membrane, αSyn has a strong binding affinity for mitochondria ^18^. Aggregated αSyn can disrupt mitochondrial membrane integrity and lead to toxicity ^19^. To examine why inhibiting the MPC results in more αSyn aggregation, we investigated if altering mitochondrial function, or changing oxidative environment via MSDC-0160 affects αSyn binding or αSyn oxidation status. Stereological quantifications of the SNpc for nitrated ni-αSyn (ni-αSyn) shows that in the αSyn low group, inhibiting the MPC with MSDC-0160 leads to more ni-αSyn (*p = 0.021, 95% CI 11.5 – 111.7 after negative binomial regression with Benjamini-Hochberg multiple testing adjustments), indicating that inhibiting the MPC leads to increased oxidation of αSyn (Fig. 4). Since it has been shown that oxidation of αSyn can affect αSyn aggregation, it raises the possibility that in this αSyn overexpression model a metabolic switch promotes the formation of insoluble assemblies of αSyn by increasing αSyn oxidation.

**Figure 4.**
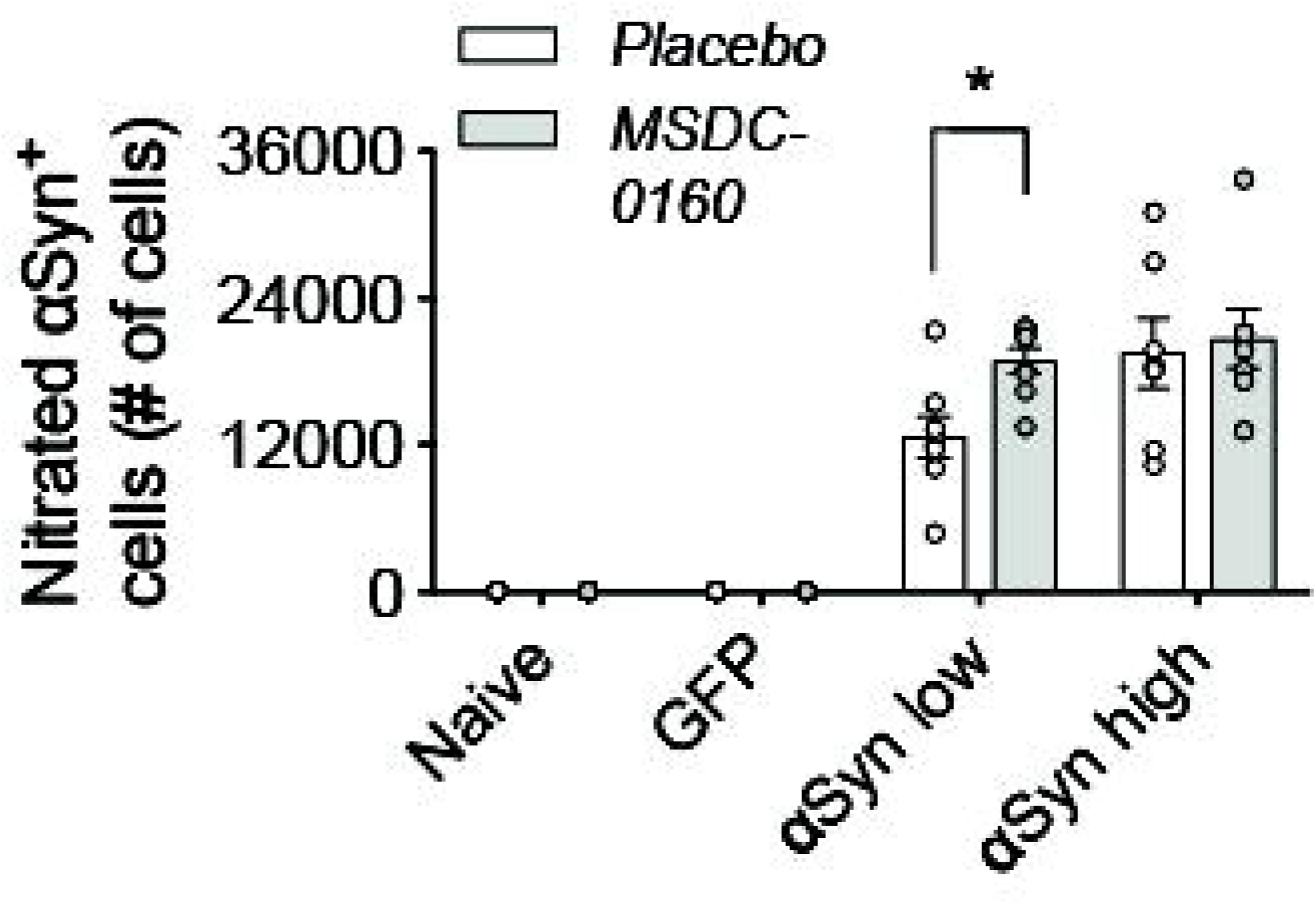
Ni-αSyn is located in the outer mitochondrial membrane and increases upon MSDC treatement. Stereological quantification of ni-αSyn positive cells in rat SN. Treatment with MSDC-0160 leads to a significant increase in nitrated cells in the αSyn low group compared to placebo (*p < 0.05 via negative binomial regression with Benjamini-Hochberg multiple testing adjustments, n = 8, SEM)

Since our previous work has shown that inhibiting the MPC has strong anti-inflammatory and metabolic effects, we further investigated if our αSyn viral vector model was lacking these features, which could explain the absence of any apparent therapeutic benefits. We examined changes in microglial and lysosomal markers at the four-month timepoint and found no changes in microglial activation (Fig. 5A-B, E) and no changes in markers of autophagy (LAMP1 and cathepsin D) in any of the groups examined (Fig. 5C, F-G). Taken together, in absence of metabolic or lysosomal deficits and microglial changes (at least at these later time points), a metabolic switch, by inhibiting the MPC, does not have a beneficial effect on αSyn aggregation.

**Figure 5.**
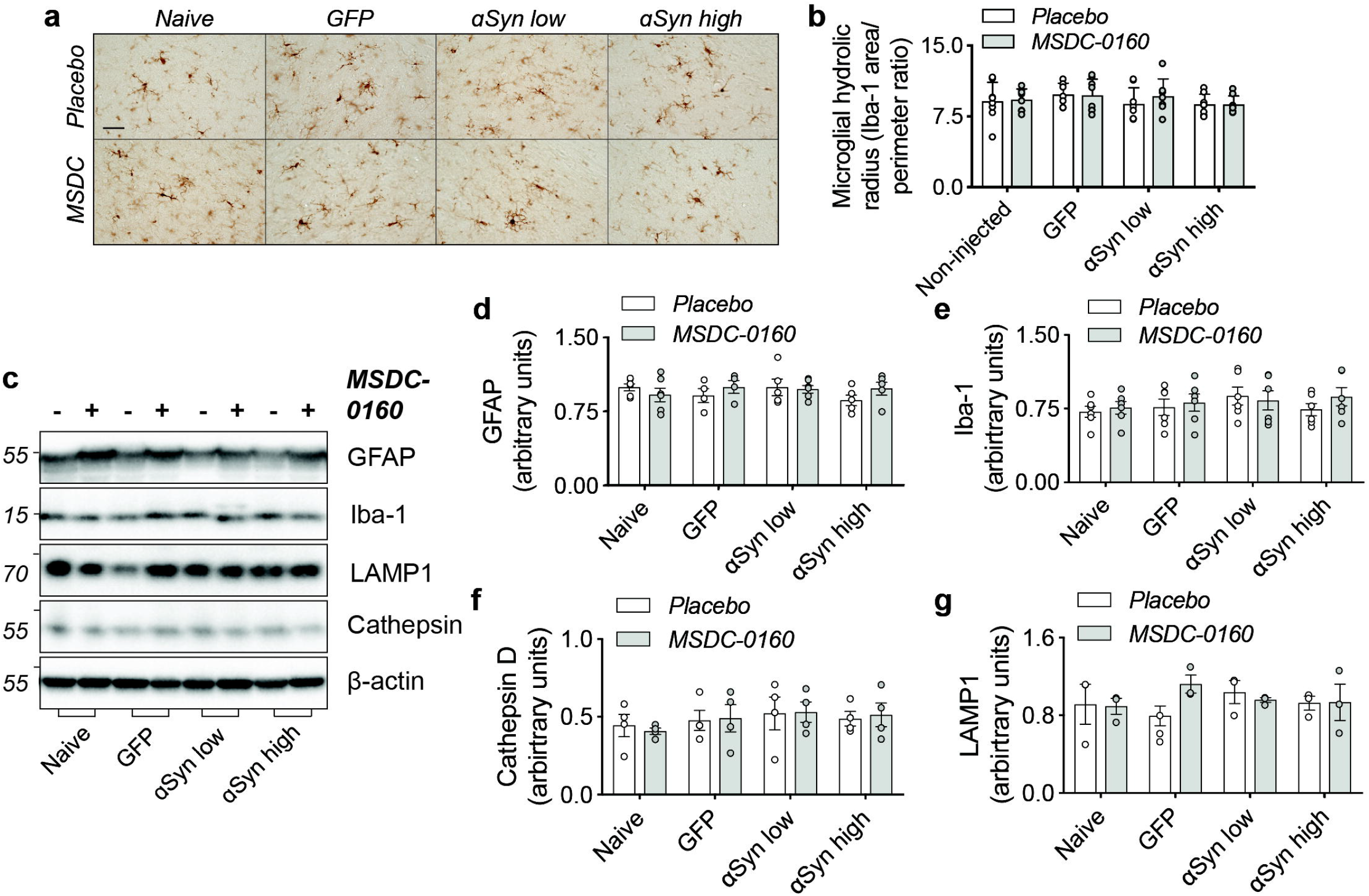
Absence of inflammatory or autophagy deficits in αSyn overexpressing animals. **a)** Overview of microglial morphology via Iba-1 staining in rat SN shows no microglial activation in αSyn overexpressing conditions. Scale bar = 50 μm. **b)** Quantification of microglial hydraulic radius as a measure of microglial ramification and activation (n = 8, SEM). **c**) Representative Western blot images of inflammatory and autophagy markers in SN. **d**) Iba-1 and **e**) GFAP markers show no signs of inflammatory activation (n = 6, SEM). **d**) Cathepsin-D and **e**) LAMP1 expression is unaltered in αSyn overexpressing animals (n = 6, SEM).

### Inhibiting the MPC in a mouse PFF seeding model

Next, to investigate the effects of inhibiting MPC on αSyn aggregation, we examined the effects of MSDC-0160 on αSyn propagation. We used a previously established seeding model where αSyn PFFs are unilaterally injected into the OB of 12-week-old wild type mice. Over time, αSyn pathology will propagate via intraneuronal connections and cause pathology in upstream regions ^19,20^. One-week post PFF or PBS injection into the OB, the mice were allowed free access to chow containing either MSDC-0160 or placebo (Fig. 6A). During the time of chow administration, we did not observe any differences in animal body weight or chow consumption between groups (Supplementary fig. 4A, B). At 5- and 13-weeks post-surgery, plasma was collected and the presence of hydroxymitoglitazone (MSDC-0160 metabolite) was quantified (Supplementary Fig. 4C), showing detectable levels of the metabolite in treated groups.

**Figure 6.**
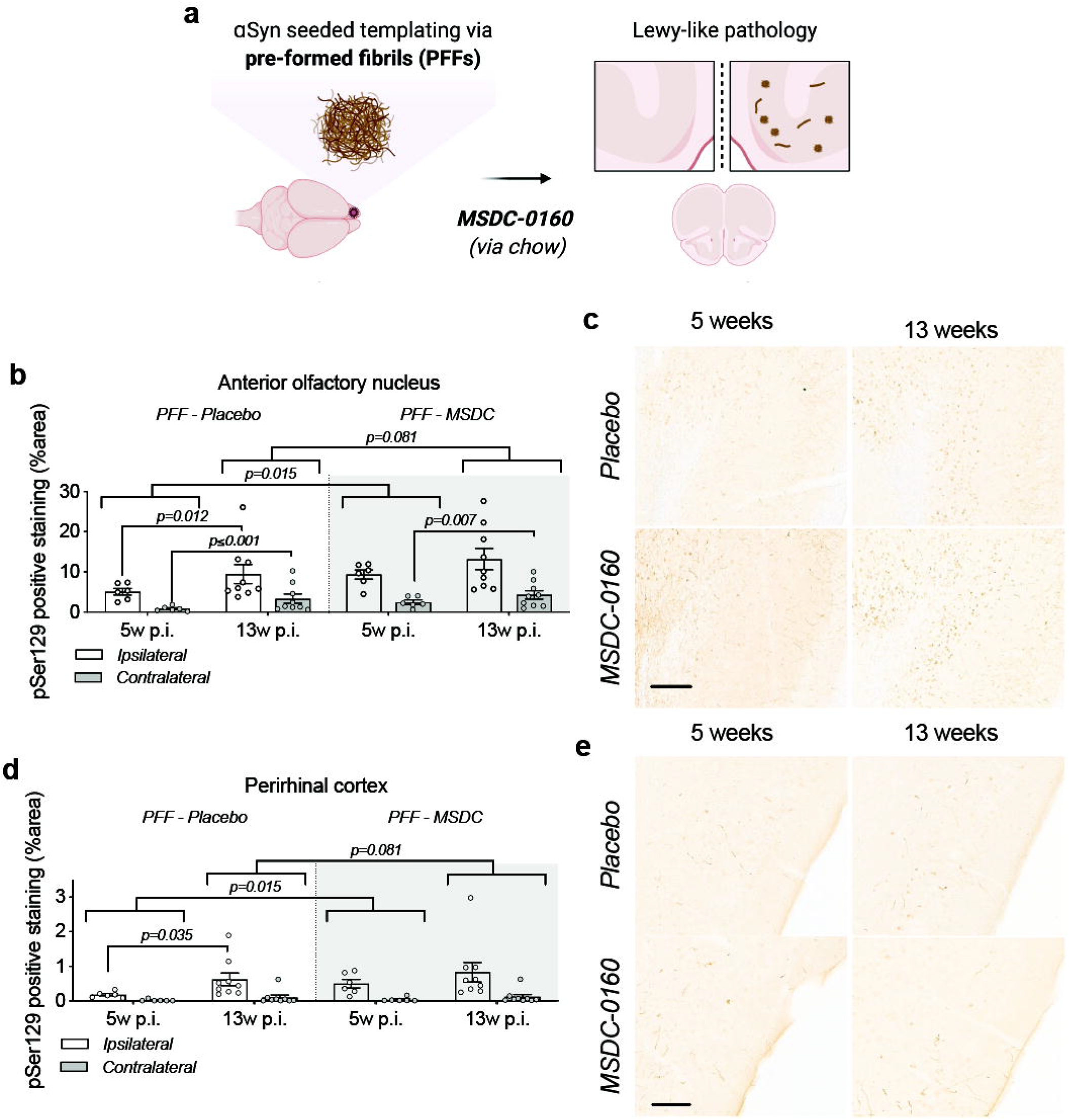
A metabolic switch increases αSyn pathology load and propagation in mouse AON and PRh. **a)** Experimental timeline; at 12 weeks of age C57BL/6J mice were injected with PFFs or PBS and starting at one-week post-surgery they had free access to MSDC-0160 or placebo containing chow until the 5- or 13-week endpoint. Quantification of αSyn pSer129 positive signal in **b)** the AON and **c)** the PRh after 5 or 13 weeks of MSDC-0160 or placebo treatment. MSDC-0160 treated mice had significantly more αSyn pSer129 positive stain at 5 weeks post-surgery than the placebo group (1.8 times more; p=0.015 95% CI 1.176 - 2.824, Beta mixed-effects regression). There is also weak evidence that this persists at the 13-week timepoint (p=0.081, 95% CI 0.986-1.92, Beta mixed-effects regression). **c)** Quantification of αSyn pSer129 positive signal in the PRh after one and three months of MSDC-0160 or placebo treatment. **d)** Representative images of αSyn pSer129 staining in **d)** the ipsilateral AON (scalebar = 130 μm) and **e)** the PRh (scalebar = 100 μm).

Unilateral PFF injections have previously been shown to impair olfactory function in mice^20^. Olfactory function was thus examined 13 weeks post-surgery using the buried pellet test. No significant difference was observed between the PBS and PFF groups, regardless of whether the mice had been treated with placebo or MSDC-0160 (Supplementary Fig. 5A). When assessing the total number of NeuN and CV-positive cells in the AON 13 weeks post-surgery via stereological quantification, we found no significant differences between any of the experimental groups (Supplementary Fig. 5B & C).

Immunohistological assessment at 5 and 13 weeks post-surgery revealed a significantly higher percentage area containing pathological αSyn (αSyn pSer129) in the AON than the PRh, regardless of time point and treatment group (9.5-18 times higher; all p ≤ 0.001) (Fig. 6B-E). The same was seen for the ipsilateral side compared to the contralateral side (67-79% less on the contralateral side; p ≤ 0.001). From 5 to 13 weeks post-surgery, αSyn pSer129 increased significantly for MSDC-0160 and placebo treated animals, which both gained ~109.6% and 177.8% of their αSyn pSer129 immunopositive stained area from week 5 to week 13, respectively in the contralateral AON (p = 0.007 and < 0.001, respectively, 95% CI = 27.7 - 243.6 and 71.2 - 350.0). In addition, placebo-treated animals had a significant increase of pathology in the ipsilateral AON from the 5 to 13-week timepoint (p=0.012, 95% CI 0.35-0.845). This was not observed for MSDC-0160 treated mice. When comparing treatment groups, MSDC-0160 treated mice had significantly more pSer129 positive stain at 5 weeks post-surgery than the placebo group (1.8 times more; p=0.015 95% CI 1.176 - 2.824). There is also weak evidence that this persists at the 13-week timepoint (p=0.081, 95% CI 0.986-1.92).

For detection of potential amyloid (ß-sheet rich) structures in the PFF seeding model and the AAV overexpression model, we used a luminescent conjugated oligothiophene (LCO). LCOs are small molecules with a flexible thiophene backbone that upon binding to amyloid structures become fluorescent^21^. One such LCO, which has previously been shown to bind aggregates in tissue samples from patients suffering from systemic amyloidosis, as well as a variety of other disease-associated protein aggregates, is h-FTAA^21^ In the AON at three months post-PFF injection, we observed ring-like h-FTAA positive structures (Fig. 7) in the PFF model. These overlapped with αSyn pSer129 antibody staining, indicating that the αSyn pathology detected in this model is ß-sheet rich with amyloid properties. On the contrary, we did not find a h-FTAA positive signal in the αSyn high group for the AAV overexpression model. This highlights differences in αSyn assembly state that results several weeks after surgery in the two PD rodent models.

**Figure 7.**
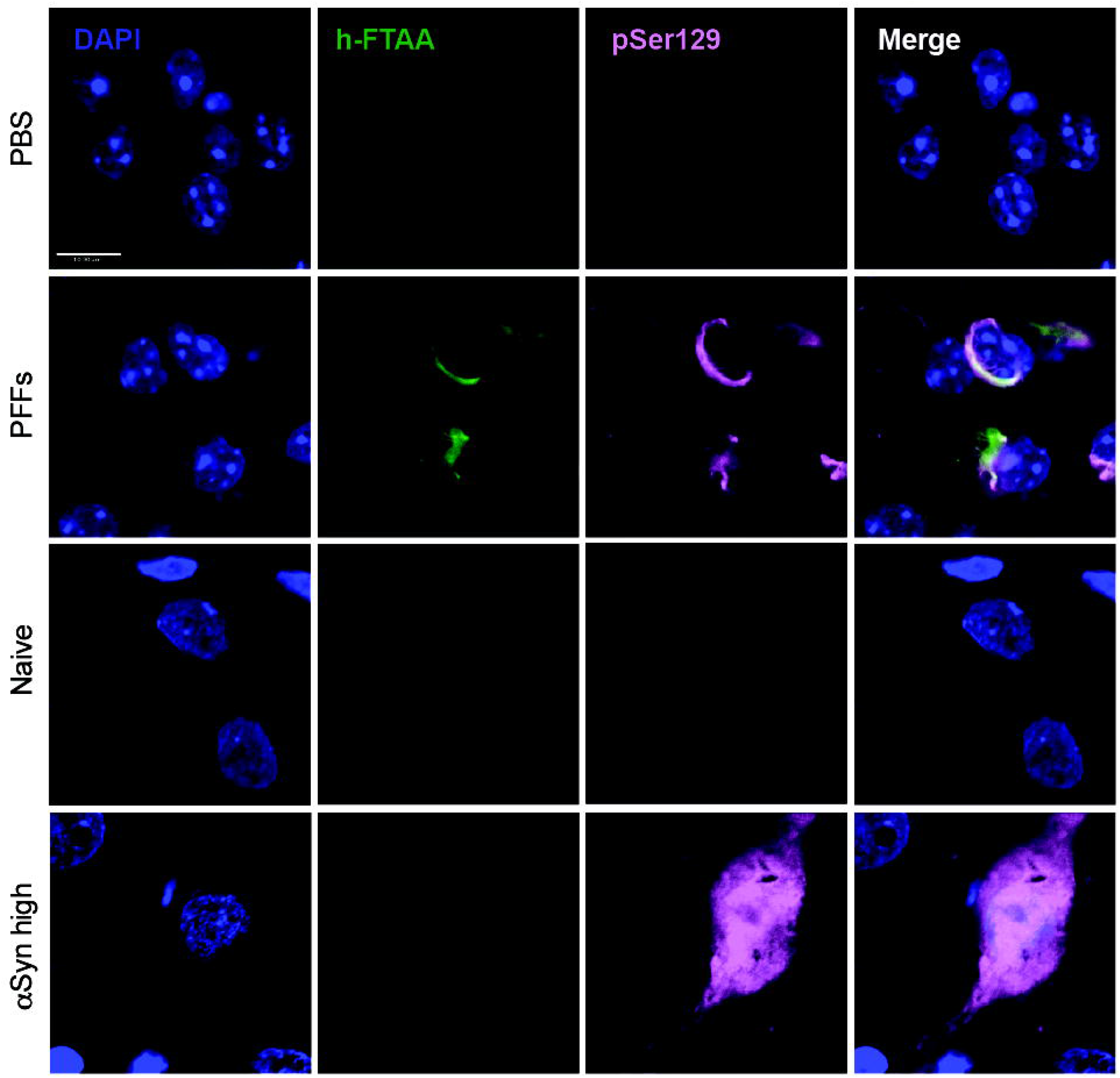
h-FTAA positive protein aggregates detected in the PFF seeding model but not the AAV overexpression model. Representative images of the AON and the SN for the PFF model and the AAV overexpression model, respectively. The h-FTAA positive signal (green) detected in the AON 13 weeks following surgery co-localizes with the αSyn pSer129 signal (pink). DAPI was used to visualize the cell nuclei (blue), scale bar = 10 μm and 3 animals per group were analyzed.

Given the absence of any therapeutic benefits in the PFF model, we again investigated the presence of inflammation via measuring the hydraulic radius (area/perimeter ratio) of microglia in the AON three months post-injection. When we compared the injected side with the non-injected side, we found no significant change in microglia morphology (Fig. 8A, B). Furthermore we did not detect any significant difference between PBS and PFFs groups or differences between animals treated with placebo or MSDC-0160. In addition, by quantifying LAMP2-positive staining we do not observe any lysosomal deficits in the AON three-months post-injection (Fig. 8C, D). Taken together, in absence of lysosomal deficits and microglial changes, a metabolic switch, by inhibiting the MPC, does not reduce accumulation of αSyn pSer129 in mice injected with PFFs in the olfactory bulb.

**Figure 8.**
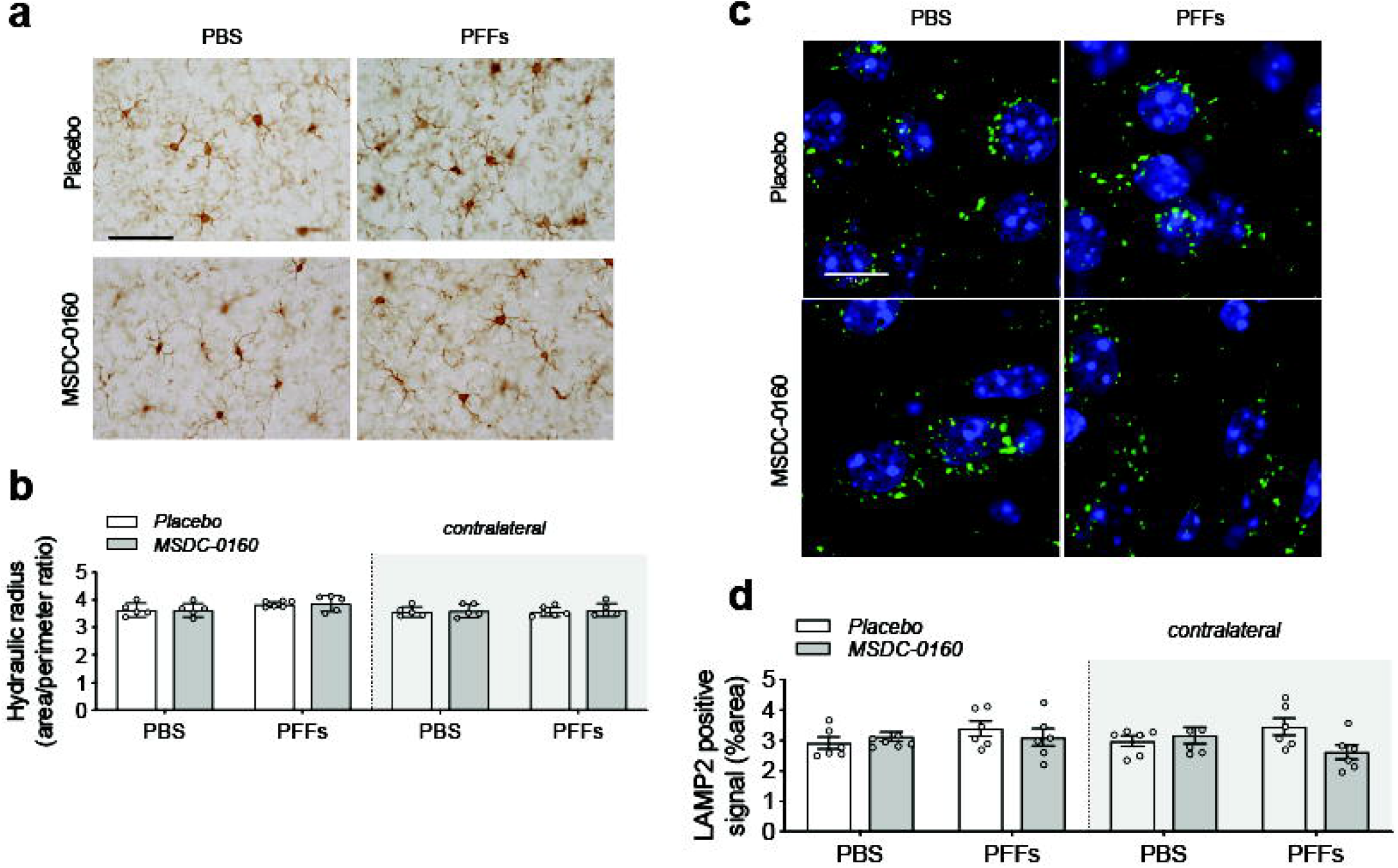
Absence of inflammatory or autophagy deficits 13-weeks p.i. **a)** Representative images of Iba-1 positive staining in the ipsilateral AON, scale bar = 50 μm. **b)** Quantification of microglial hydraulic radius as a measure of microglial ramification and activation (n = 5-6/group, SEM), shows no microglial activation following PFF injection. **c)** Representative images of LAMP2 positive staining (green) in the ipsilateral AON. DAPI (blue) was used to visualize the cell nuclei. Scale bar = 10 μm **d)** Fluorescent LAMP2 quantification after adding adaptive triangle thresholding in the AON (n = 6, SEM)

## Discussion

Improving metabolism, reducing inflammation and inhibiting the aggregation or the spread of pathological αSyn are considered a promising therapeutic targets for the treatment of PD. Through preclinical and clinical work, it has been suggested that anti-diabetic drugs might hold this promise^22^. We have previously shown that MSDC-0160, a type 2 diabetes insulin sensitizer, can ameliorate pathology in an acute (1-methyl-4-phenyl-1,2,3,6-tetrahydropyridine, MPTP) and slowly progressive, genetic (Engrailed1^+/-^) model of PD and that these effects occurred via modulation of the MPC and its downstream effect involving the mTOR pathway^10^. Since the direct effects of inhibiting MPC in chronic models of αSyn aggregation had not been studied, we asked if MSDC-0160 could lower the levels of αSyn in two different, chronic and progressive αSyn-based animal models of PD. In our first model, αSyn is expressed via viral vector delivery in the rat SN resulting in aggregation and progressive accumulation of insoluble αSyn. After four months, no significant motor abnormalities with moderate neurodegeneration was observed but, unexpectedly, animals that were treated with MSDC-0160 exhibited increased levels of aggregated αSyn. In the second model, PFFs were injected in the OB of mice and allowed to seed and spread to connected regions. Five weeks after PFF injection, we observed an increase in αSyn pathology burden when mice were treated with MSDC-0160. This effect was transient and at the later, 13-week timepoint no significant difference between the two treatment groups could be observed.

Both the αSyn viral vector and the αSyn PFF seeding models are based on the introduction of exogenous αSyn that leads to progressive accumulation of phosphorylated and misfolded αSyn over time. The viral vector model results in local accumulation of highly concentrated but soluble αSyn that assembles into insoluble αSyn, whereas the PFF model is based on the introduction of aggregated, seeding-potent αSyn that relies on the endogenous protein to template and spread via anatomically connected regions. To date, there are no studies that have directly compared the assembly states of the αSyn in the intraneuronal inclusions that remain after several weeks in an overexpression versus a seeding model, but it appears that inhibiting the MPC has different outcomes in these two models. This tentatively suggests i) that the conformational state or aggregation state of αSyn might be different in the two models and ii) that the effects of inhibiting the MPC depends on the assembly state of the protein.

We observed a disparity in αSyn assembly states between the viral vector-based and the seeding model in multiple assays. First, we used detection of nitrated αSyn to determine that αSyn is oxidized in the viral vector-based model. A previous study showed that αSyn can bind directly to the mitochondrial import protein TOM20, which leads to increased mitochondrial respiration and ROS production^17^. We now find that oxidized αSyn species are abundant in neurons in the viral vector model, but undetectable in the PFF model. We postulate that by altering the TCA cycle when treating mice overexpressing αSyn with MSDC-0160, the oxidation of αSyn is increased. This suggests that alterations in mitochondrial metabolism might post-translationally modify αSyn and make it more aggregation prone, when the protein is highly concentrated. Furthermore, we propose that elevated levels of soluble αSyn are required for these changes to take place, since we did not observe similar effects in the PFF seeding model where the endogenous levels of αSyn are normal and aggregates are formed via an induced seeding mechanism. Since the conformational state of αSyn is closely linked to its function^23^, it is possible that elevated levels of monomeric αSyn lead to changes in assembly state or gain of post-translational modifications resulting in an aberrant interaction with mitochondria. The binding affinity of αSyn is higher for mitochondrial membranes due to the unique presence of cardiolipin in mitochondria^18^. Lipid interaction is an important catalyst for soluble αSyn to convert into aggregated αSyn^24^ and this conversion has been reported to lead to release of mitochondrial nitric oxide and increased oxidative stress^25^. αSyn can disrupt membrane integrity and cause mitochondrial damage and degeneration of dopaminergic neurons^17,26^. Recombinant nitrated species of αSyn injected in rat SN have been shown to be more toxic than unmodified αSyn^27^. In MSDC-0160 treated mice, we did indeed observe greater loss of dopamine neurons in the anterior portion of the SN, which was the region where viral vector-mediated expression of αSyn was the highest.

Multiple molecules of nitrated αSyn have been shown to form soluble assemblies that are off pathway to aggregation and relatively unstructured^28^. In order to determine if structured amyloid forms of αSyn were present in our animal models, we used the conformation sensitive LCO h-FTAA probe^29^. LCOs are amyloid-conformation-specific dyes that specifically bind to amyloid αSyn in brain tissue and CSF^30^. A positive signal therefore indicates the presence of well-structured, ß-sheet rich aggregated αSyn. In addition, h-FTAA has been used to distinguish protein inclusion bodies in sporadic inclusion body myositis (s-IBM) and liver diseases^21^. Upon treatment with h-FTAA, we observed a strong positive signal in the AON of animals injected with PFFs but no signal in the SNpc of animals overexpressing αSyn following viral vector injection. The fluorescent signal was located to neurons and the morphology was Lewy body-like with aggresomal structures that resembled the intracellular distribution we observed when staining for αSyn pSer129. This illustrates that the two αSyn animal models also differ in that the αSyn assembly states in these models are distinct. These differences were recently discussed by Gómez-Benito and colleagues; αSyn species from the AAV overexpression model lack seeding activity while the PFF model recapitulates αSyn propagation with the accumulation of Lewy body-like structures^31^.

In summary, MSDC-0160 does not reduce αSyn aggregation in the model where a viral vector is used to overexpress the protein in the nigrostriatal system. On the contrary, the inhibition of MPC results in a slight increase in αSyn protein content. In the model using injection of PFFs to trigger a cascade of αSyn aggregate propagation in the olfactory system, MSDC-0160 exhibits no effect on the progressive development of synucleinopathy in the olfactory system. In experimental paradigms that involve more acute neuroinflammation, we previously showed that MSDC-0160 alleviates the inflammatory response *in vitro* and *in vivo*^10^. However, in the PFF and viral vector αSyn models, we did not detect changes in inflammation or evidence for autophagy deficits at the survival times we studied. We conclude that attenuation of the MPC rodent models that do not exhibit inflammation and limited autophagy does not result in a reduction of introduced synuclein.

## Acknowledgements

Research reported in this publication was supported by the Cure Parkinson’s trust award number 42-40384-1 (P.B). WB acknowledges a post-doctoral fellowship from Fulbright, IDT Technologies and FWO Flanders. We thank the staff of the Vivarium of Van Andel Institute for caring for the mice and rats used in this study.

## Author contributions

W.P performed rat studies (stereotaxic surgery and behavioral, biochemical and immunohistochemical analysis) and wrote the manuscript. L.B performed mouse olfactory testing, NeuN staining and stereology, pSer129 staining and analysis, ni-αSyn staining, LAMP2 staining and analysis, Iba-1 staining and analysis, h-FTAA/pSer129 co-staining and imaging and wrote the manuscript. S.G. performed pilot studies leading to the final conceptualization of the AAV overexpression model study. M.E.J performed pSer129 staining. J.S organized the manufacturing of the MSDC-0160 chow and provided advise. L.M performed unilateral PFF injections and mouse perfusions. E.K performed unilateral PFF injections, mouse perfusions and ni-αSyn staining optimization. Z.M. performed the statistical analysis. J.M provided the PFFs. K.B performed PFF manufacturing. J.C provided the compound (MSDC-0160), expertise and advice. P.B. conceptualized this study and provided expertise and advice. All authors reviewed and edited the manuscript.

---

**Supplementary Figure 1. Body weight, chow consumption and plasma MSDC-0160 metabolites**. **a)** Body weight of female rats during 16 weeks of MSDC administration (n = 80, s.e.m.) **b**) Chow consumption of placebo or MSDC-0160 infused chow. Intake was averaged over two weeks per animal (n = 20, s.e.m.) **c**) Plasma levels of hydroxymitoglitazone, the active metabolite of MSDC-0160 in female rats, 16 weeks after start of MSDC-0160 administration. No metabolites were detected in placebo treated animals (n = 3, s.e.m.).

**Supplementary Figure 2. Motor behavior analysis of MSDC-0160 treated animals.** Cylinder test for αSyn overexpressing and control animals shows no significant changes or motor deficits in animals treated with MSDC-0160 (n = 40, s.e.m.).

**Supplementary Figure 3. Nigral cell death anterior in SN. a)** Fluorescent detection of TH-positive dopaminergic neurons in the ipsilateral anterior rat SN. Binarized images were thresholded by the adaptive triangle method in ImageJ. Scale bar indicates 500 μm. **b)** Quantification of TH positive area after thresholding shows significant cell loss in the anterior part of rat SN (n = 8, s.e.m., #p < 0.05 for two-way ANOVA with Tukey post hoc correction compared to naïve condition treated with MSDC-0160). No differences between treatment conditions were detected. **c)** Fluorescent TH detection after adaptive triangle thresholding of ipsilateral posterior rat SN. Scale bar indicates 500 μm. **d)** Quantification of TH positive area after thresholding shows no cell loss in the posterior part of rat SN (n = 8, s.e.m.).

**Supplementary Figure 4. Absence of inflammatory or autophagy deficits in αSyn overexpressing animals. a)** Overview of microglial morphology via Iba-1 staining in rat SN shows no microglial activation in αSyn overexpressing conditions. Scale bar indicates 50 μm. **b)** Quantification of microglial hydraulic radius as a measure of microglial ramification and activation (n = 8, s.e.m.). **c**) Representative Western blot images of inflammatory and autophagy markers in SN. **d**) Iba-1 and **e**) GFAP markers show no signs of inflammatory activation (n = 6, s.e.m.). **d**) Cathepsin-D or **e**) LAMP1 expression is unaltered in αSyn overexpressing animals (n = 6, s.e.m.).

