## Supplementary figures and images for "Inhibiting the mitochondrial pyruvate carrier does not ameliorate synucleinopathy in the absence of inflammation or metabolic deficits"

### Supplementary figure 1

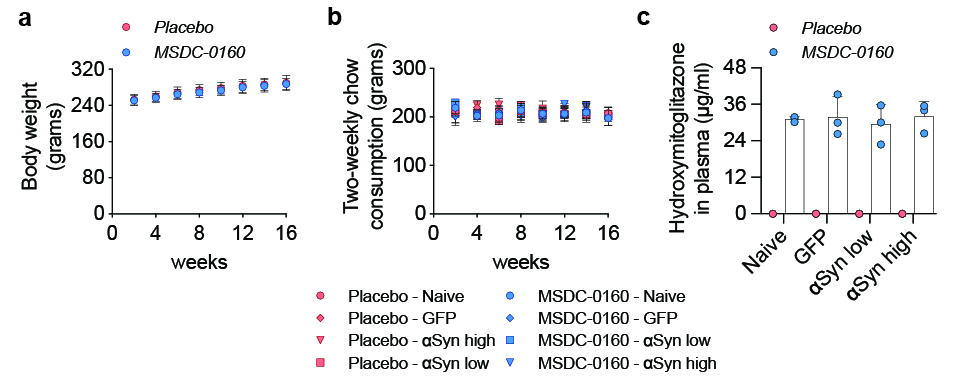

### Supplementary figure 3

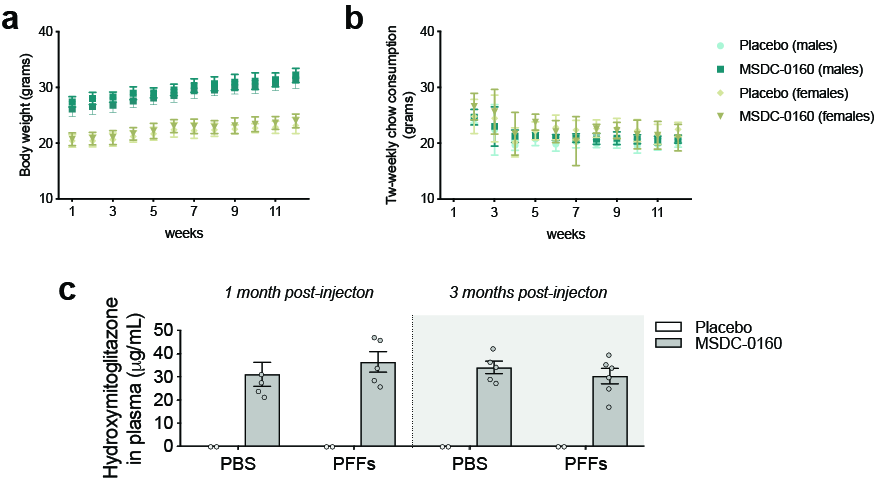

### Supplementary figure 4

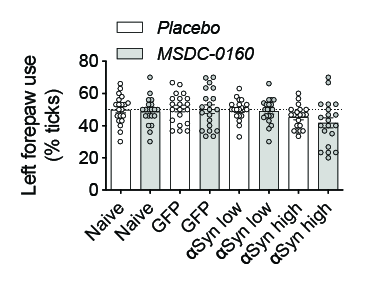

### Supplementary figure 4

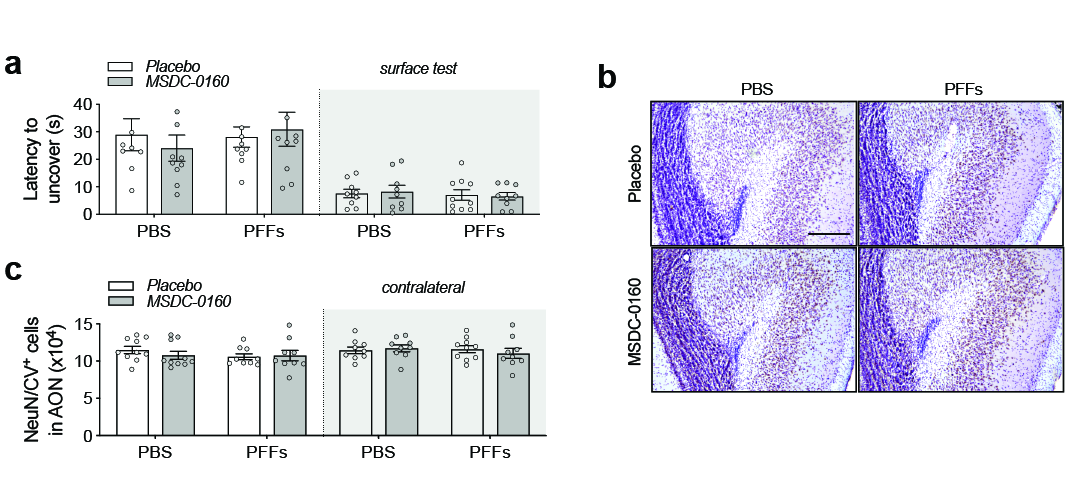

### Supplementary figure 5

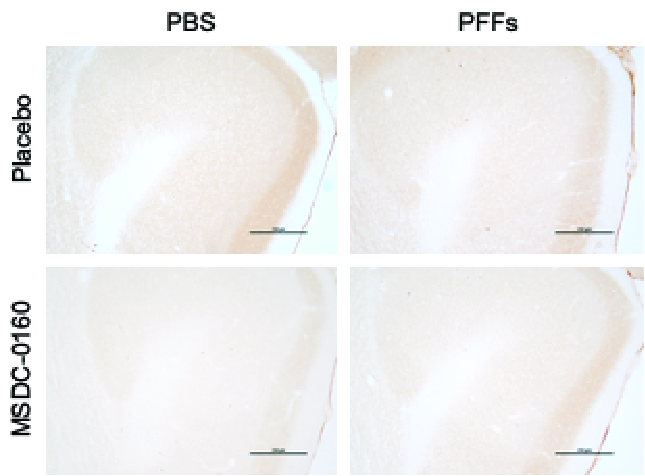
